# A Spatial Transcriptomics based Label-Free Method for Assessment of Human Stem Cell Distribution and Effects in a Mouse Model of Lung Fibrosis

**DOI:** 10.1101/2023.05.31.542821

**Authors:** Jeongbin Park, Dongjoo Lee, Jae Eun Lee, Daeseung Lee, In Ho Song, Hyun Soo Park, Hongyoon Choi, Hyung-Jun Im

## Abstract

Recently, cell therapy has emerged as a promising treatment option for various disorders. Given the intricate mechanisms of action (MOA) and heterogenous distribution in target tissues inherent to cell therapy, it is necessary to develop more sophisticated, unbiased approaches to evaluate the distribution of administered cells and the molecular changes at a microscopic level. In this study, we present a label-free approach for assessing the tissue distribution of administered human mesenchymal stem cells (hMSCs) and their MOA, leveraging spatially resolved transcriptomics (ST) analysis. We administered hMSCs to mouse model of lung fibrosis and utilized ST to visualize the spatial distribution of hMSCs within the tissue. This was achieved by capitalizing on interspecies transcript differences between human and mouse. Furthermore, we could examine molecular changes associated with the spatial distribution of hMSCs. We suggest that our method has the potential to serve as an effective tool for various cell-based therapeutic agents.

## Introduction

Cell therapy involves the introduction of therapeutic cells into patients and has gained significant attention as a promising treatment option for a range of diseases. It has garnered significant attention in recent years as a promising treatment modality for various diseases, including cancers [1], autoimmune diseases [2], inflammatory diseases [3], and neurodegenerative diseases [4]. Cell therapeutic agents are characterized by designability, biocompatibility, and applicability of cell functions even surpassing blood-brain barrier (BBB) [5], making cell therapy more promising. With the advent of advanced techniques in cellular manipulation (*e*.*g*., chimeric antigen receptor-T cell or NK cell therapy (CAR-T/NK), stem cells, tumor infiltrating lymphocytes, or microbiome agents), the potential of cell therapy as a game-changing therapeutic approach continues to grow. Despite the promise of cell therapy, it has faced significant challenges due to the living nature of cells, resulting in limitations in reproducibility of drug efficacy and difficulties in preparation, delivery, and administration [6].

Traditional methods assessing the efficacy of cell therapy often lack the necessary resolution and accuracy to provide meaningful insights into the complex interactions and molecular changes that occur during treatment. For example, poly-chain reaction (PCR) is basically performed in organ resolution, limiting the identification of the microscopic properties of cell therapy, particularly in-tissue distribution of administered cells [7]. Fluorescence-labeled cells, which are commonly used to obtain images of administered cells, have a drawback in that the tracers can easily detach from the cells, resulting in limitation for tracking cells [8]. Most of all, it is important to note that these methods do not provide a comprehensive understanding of the detailed molecular changes and interactions that occur between the administered cells and the host cells within the target tissues. These have led to a pressing need for innovative and comprehensive approaches to better understand the true potential of cell therapeutic agents.

Spatially resolved transcriptomics (ST) has emerged as a groundbreaking tool in the field of molecular biology [9]. By combining spatial information with gene expression data, this technique allows researchers to study the dynamic changes in gene expression patterns within tissues and cells at an unprecedented level of detail. ST has the potential to revolutionize the evaluation of cell therapeutic agents by providing a more comprehensive understanding of their effects on the cellular and molecular level.

In this paper, we present a novel experimental and analytic procedure that harnesses the power of ST to evaluate cell therapeutic agents. This approach not only offers a more in-depth understanding of the mechanisms underlying the therapeutic effects of these agents, but also provides critical insights into the optimization of cell therapy treatments for various diseases. Here, we administered human mesenchymal stem cells (hMSC) in a mouse model of lung fibrosis and presented a method for evaluating both the tissue-level distribution of the administered stem cells and their effect on treated tissue. We expect this analysis method to facilitate the development of cell therapeutics by comprehensively understanding mode of action and in-depth cell-level distribution in the microenvironment.

## Results

### Detection of human transcripts for label-free evaluation of stem cell distribution

The overall research design and ST library preparation for lung tissues from normal mouse (‘*Nor*’), lung fibrosis model (‘*Con*’), and lung fibrosis model treated with hMSC cells (‘*Exp*’) were represented in **Figure 1a**. Lung fibrosis was induced by intravenous injection of bleomycin for 3 weeks. Lung fibrosis was visually identified on histopathologic images of lungs (**Supplementary Figure 1**). Normal bone marrow derived hMSC was intravenously injected in the ‘*Exp*’ mouse 6 h after sacrifice. The cDNA library of spatially resolved transcriptomics (ST) was mapped using a reference genome that combined both human and mouse genomes to detect human transcripts from the administered cells. With constant sequencing depth, the number of raw counts of a gene in a single spot is inversely proportional to the total RNA production for that spot. To address this, gene counts are adjusted using various methods, such as the trimmed mean of M-values (TMM) [10] and fractile normalization. Also, since genes with large counts can overestimate their activities, gene counts are generally log-transformed. This process, referred to as normalization, can be performed using *scanpy*.*pp*.*normalize_total* and *scanpy*.*pp*.*log1p* functions in Python. Following the normalization of gene counts, the distribution of all the transcripts became more closely aligned with normal distribution (**Supplementary Figure 2**, the first column). The percentage of the human transcripts of all the transcripts for a single spot, %human, was also explored according to sample and normalization. As a result, %human with normalization was higher in the ‘*Exp*’ sample compared to without normalization (**Supplementary Figure 2**, the right bottom), while having minimal impact on the ‘*Nor*’ and ‘*Con*’ samples. Hence, we focused on human transcripts and %human with normalization for the subsequent analysis. We tested various thresholds to discriminate falsely detected human transcripts (**Supplementary Figure 3**). Consequently, we defined a threshold based on the average plus 10 times the standard deviation of %human in the ‘*Con*’ sample for two reasons. Firstly, this threshold was just above the critical value of %human where no spots exceeding it in the ‘*Nor*’ and ‘*Exp*’ samples (**Supplementary Figure 2**, the third column). Secondly, when applying this threshold, the classified spots with human cells in the ‘*Exp*’ sample displayed a histological pattern suitable for comparison in the following cell type matching analysis.

**Figure 1.**
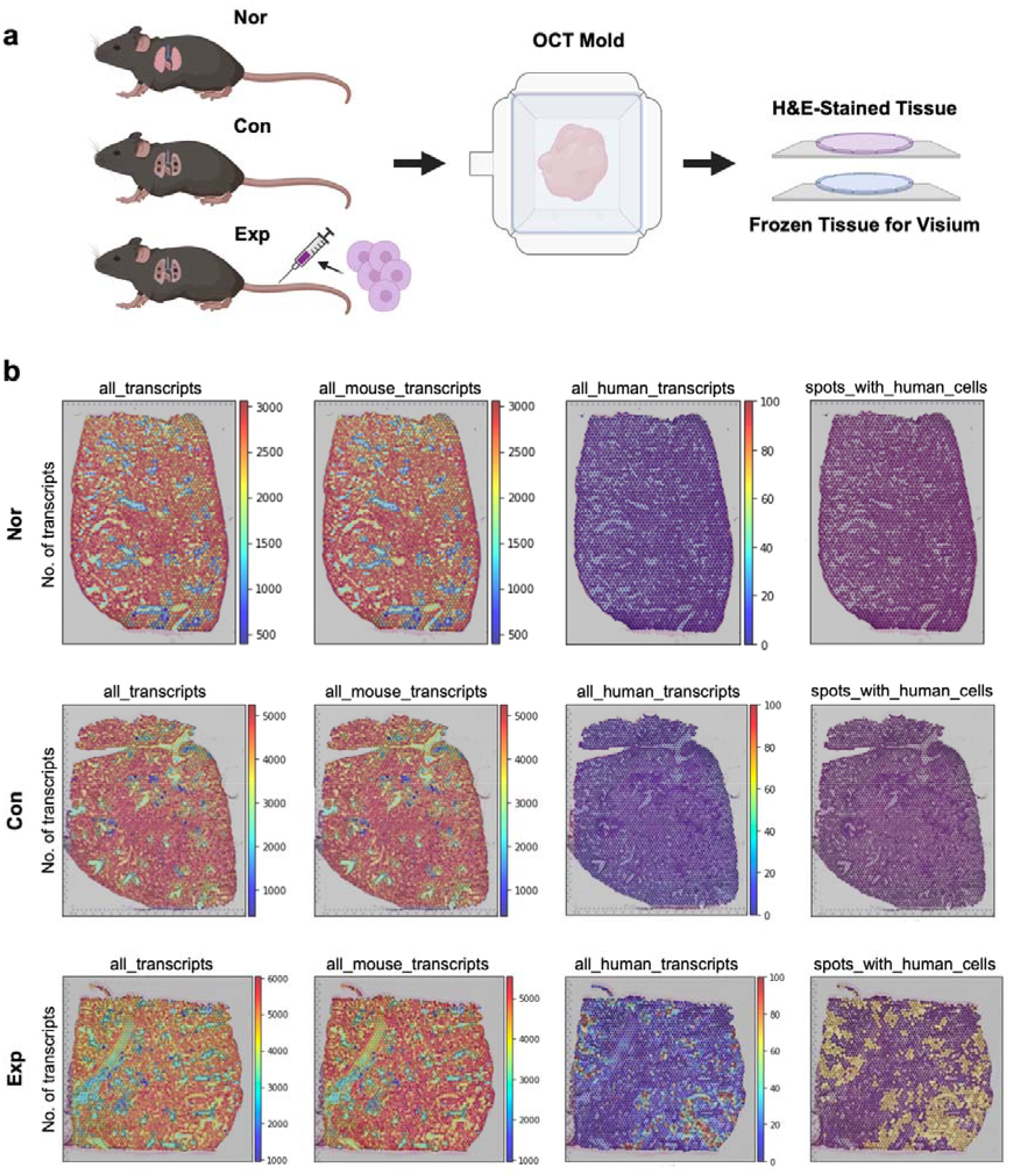
Preparation of lung tissues from normal mouse (‘*Nor*’), lung fibrosis model (‘*Con*’), and lung fibrosis model treated with hMSC cells (‘*Exp*’). **a**, Schematic illustration of ST library preparation. Lung fibrosis and hMSC injection were also represented. **b**, Number of transcripts according to sample and organism. Here, transcripts indicated not raw gene counts, but normalized gene counts. The results for ‘*Nor*’, ‘*Con*’, and ‘*Exp*’ were represented from top to bottom. The last column of the image, ‘*spots_with_human_cells*’, showed a binary indicator that was yellow only when %human on a spot is greater than the average plus 10 standard deviations of %human of the ‘*Con*’ sample. Spatial mapping of ‘*spots_with_human_cells*’ well represented the existence of injected human stem cells in the ‘*Exp*’ sample.

The number of human transcripts was only observable in ‘*Exp*’ sample compared to the others (**Figure 1b**, scale of 0-100 transcript number, the third column), although low number of human transcripts were also found in the ‘*Nor*’ and ‘*Con*’ samples (**Supplementary Figure 2**). Given that the ‘*Nor*’ and ‘*Con*’ samples were not administered human cells, the human transcripts detected in the samples could be considered as false positives. However, as the false positive thresholding was effective, the distribution of %human on ST could be considered as distribution of administered hMSC (**Figure 1c**, the fourth column).

### Clustering analysis identified administered hMSCs in the lung tissue

As a result of spatial clustering analysis based on mouse gene expression, 9 clusters were identified according to the gene expression patterns (**Figure 2a**). Among them, cluster 5 showed significantly higher %human than others (**Figure 2b**). In addition, spots of the cluster 5 were rarely appeared in ‘*Nor*’ and ‘*Con*’ samples (**Figure 2c**). It was remarkable that even if human genes were excluded when performing spatial clustering analysis, significantly higher number of human transcripts were concentrated on a single cluster. Top 20 spatially enriched mouse genes in the cluster 5 implied the occurrence of immune response, collagen-containing extracellular matrix (ECM), and peptidase activity (**Figure 2d**). Differentially expressed genes (DEGs) between ‘*Con*’ and ‘*Exp*’ were obtained. Although we included all genes from human and mouse, top 6 DEGs were all mouse genes (**Supplementary Figure 4-6**). To further comprehend genes spatially associated with the hMSC distribution, the spatially associated genes with %human were explored in the ‘*Exp*’ sample (**Figure 3a and 3b** for positively and negatively correlative genes, respectively). In this analysis, six genes were human genes among top 20 associated genes. To assess molecular process of lung fibrosis in the ‘*Exp*’ sample, we selected positively spatially correlated mouse genes in the top 20 spatially associated genes and performed GO analysis with them. The top biological pathway was ‘*transmembrane receptor protein serine/threonine kinase signaling pathway*’ (**Figure 3c**). Also, there were overlapping mouse genes between top 20 spatially enriched genes in the spatial cluster 5 and the spatially associated genes: *Lcn2* and *Col4a1* (**Figure 2-3**).

**Figure 2.**
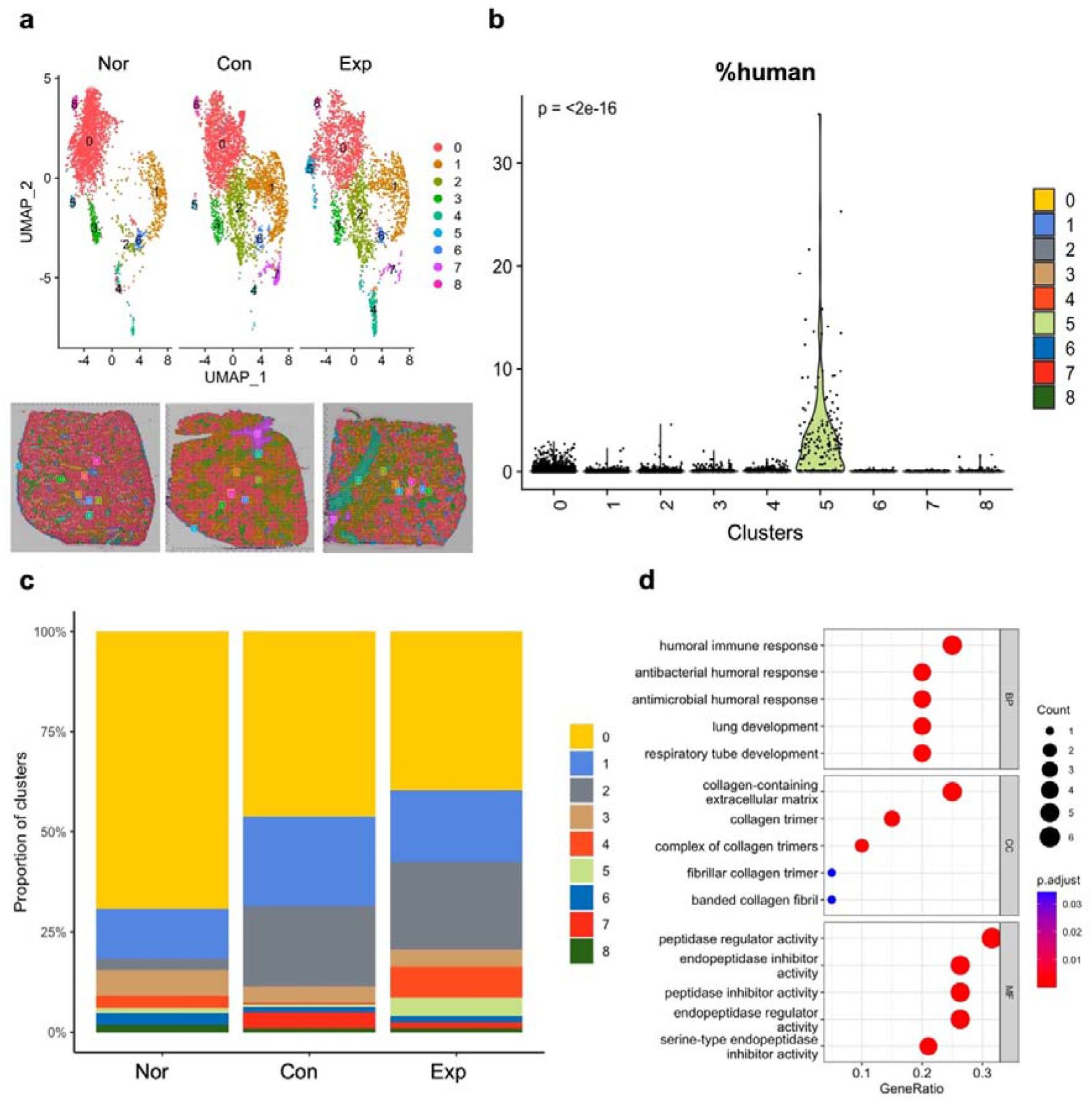
Spatial clustering analysis. **a**, *DimPlot* (*up*) and *SpatialDimplot* (*down*) of three samples according to clustering labels. **b**, *VlnPlot* for %human according to clustering labels. Interestingly, cluster 5 showed a significantly higher number of human transcripts than the others. **c**, The population of clustering labels for each sample. The cluster 5 rarely appeared in ‘*Nor*’ and ‘*Con*’ samples. **d**, GO plot for top 20 spatially enriched genes in the cluster 5 (adjuster p value < 0.05; log FC ordered) including *Lcn2* (highest log FC), *Msln, Chil3, C3, Upk3b, Spp1, Col4a1, Serpina3n, Wfdc21, Col3a1, Wfdc17, Gm13889, Chil1, Mgp, Sftpd, Slpi, Ctsc, Fmo2, Scd1*, and *Napsa* (in order).

**Figure 3.**
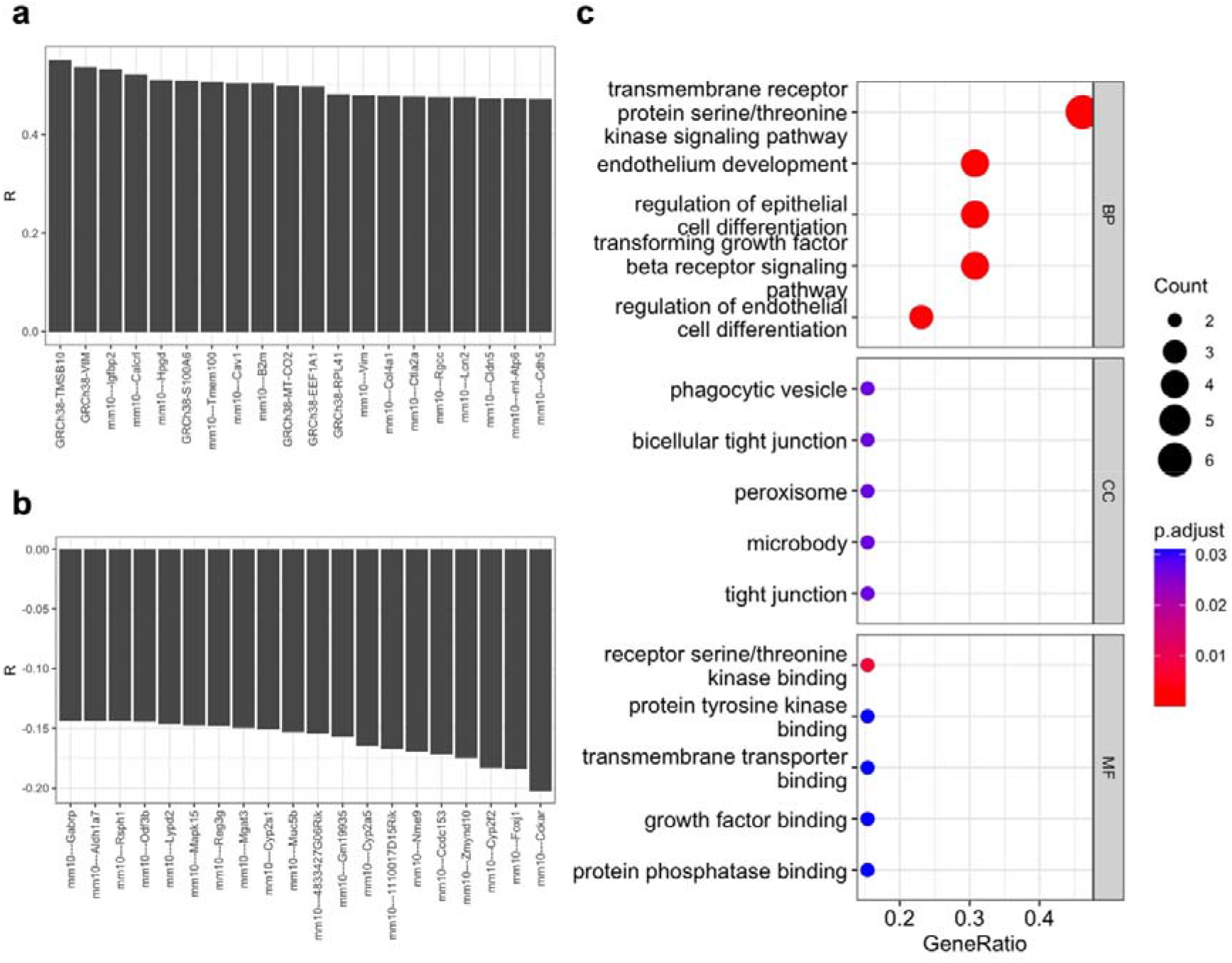
Spatially associated genes with %human in the ‘*Exp*’ sample. The most (**a**) and the least (**b**) 20 spatially associated genes to %human. **c**, GO analysis for the mouse genes among top 20 spatially associated genes to %human.

### Cell type population of lung fibrosis model associated with hMSC treatment

After preparing a single cell RNA sequencing (scRNA-seq) reference of mouse lung (GSE124872), cellular level deconvolution was performed on the ‘*Exp*’ sample using CellDART [11] (**Figure 4a**). Spearman correlation between human transcripts and cell types of the treated lung tissue was calculated (**Figure 4b**). As a result, endothelial cells and epithelial cells were the most and the least associated cell type with %human, respectively (**Fib. 4b**). It was well concurred with the fact that the migration of injected cells into lungs were thought to be physically trapped in narrow micro-vessels of the alveoli [12].

**Figure 4.**
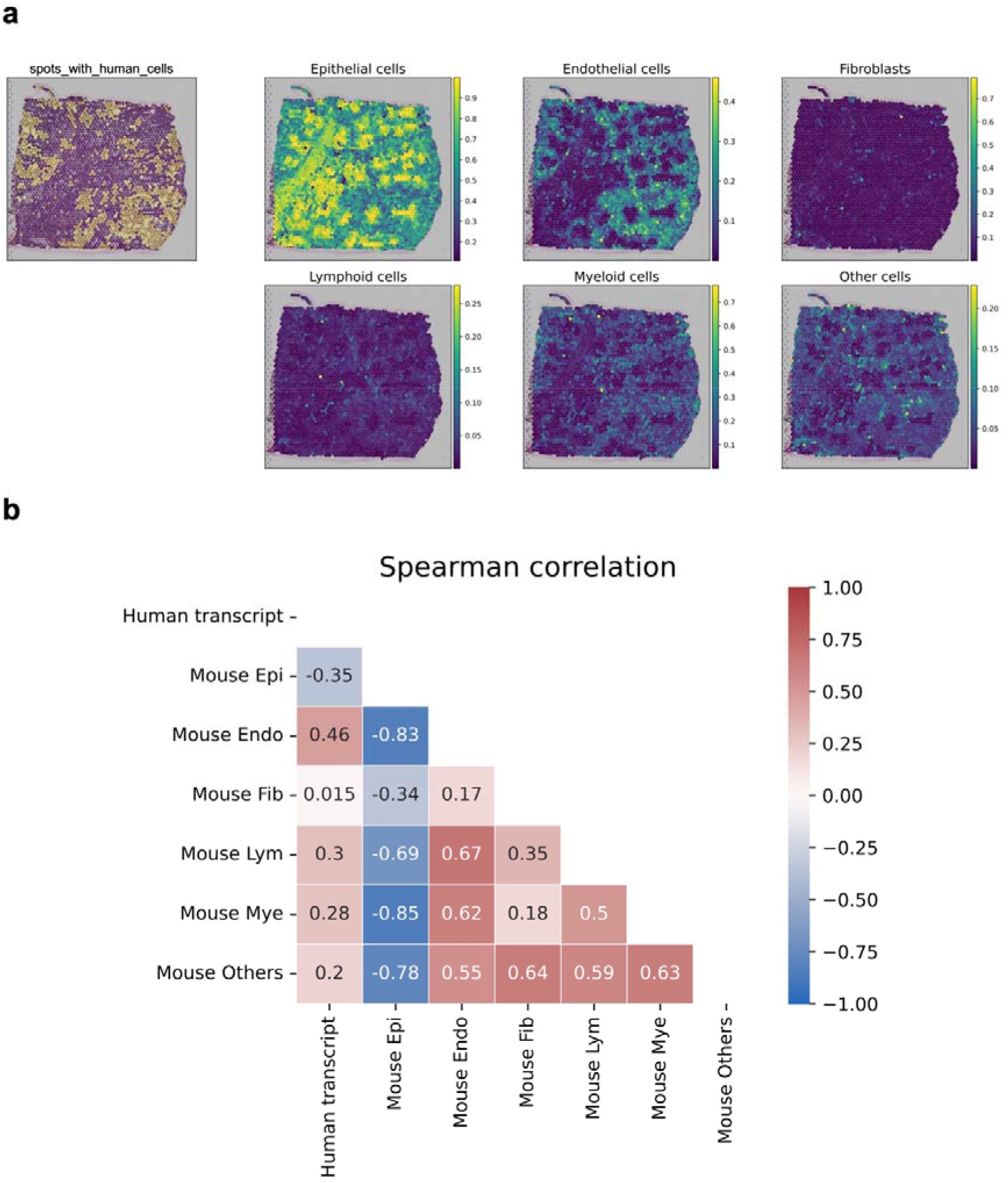
CellDART results. **a**, Spatial mapping of the proportion of 6 cell types in the ‘*Exp*’ sample, along with human transcripts representation. **b**, A correlation coefficient matrix comparing the cell type proportions, along with the normalized %human. Here, spearman correlation coefficient was used. As a result, endothelial cells and epithelial cells were the most and the least associated cell type with %human, respectively.

We defined the ‘*is_endothelial*’ variable to indicate whether a spot consists of endothelial cells in the ‘*Exp*’ sample, using a threshold of the average plus one standard deviation of the CellDART score for endothelial cells. We then obtained DEGs in the same manner as the prior DEG analysis, comparing ‘i*s_endothelial*’ spots with and without the ‘*is_human*’ designation. As a result, only *Apoe, Col1a1*, and *Hnrnpab* significantly appeared (adjusted p-value < 0.05) in spots with low human transcripts. Especially, *Apoe* and *Col1a1* (activated fibroblast marker) genes are known to be situated in stromal regions surrounding blood vessels [9b]. In contrast, *mt-Co2, Lcn2, Gm42418, Chil1, Scd1, mt-Nd1*, and *mt-Nd2*, along with 43 human genes, were significantly present (adjusted p-value < 0.05) in the spots with high human transcripts. Interestingly, *Lcn2* found in the former two analyses also appeared in this analysis, supporting the overall conclusion.

## Discussion

In this paper, we could find the following three discoveries. First, RNA transcripts originating from different genomes could be discriminated and the spatial distribution of them could be acquired. Second, significantly associated genes with the human RNA transcripts can be obtained by spatial clustering analysis, DEG analysis, and spatially associated genes with %human. Lastly, the injected human stem cells displayed a strong association with endothelial cells, while showing an inverse association with epithelial cells.

The market for therapeutic drugs containing nucleic acids is expanding, with numerous drugs currently in development. Clinical trials for stem cells [13], CAR-T [14], and exosomes containing nucleic acids [15] have been conducted. Nonetheless, there have been reports of challenges in evaluating the molecular mechanisms of these drugs particularly interacting with cells in target tissues in preclinical studies. As a result, there is an urgent need for the assessment methods for these drugs. Earlier, a method to identify molecular markers spatially associated with an injected drug based on ST was developed [9b]. Using this method, markers related to enhanced permeability and retention (EPR) were identified and it can be applied to various therapeutic agents that are labeled with fluorescent dyes. Nonetheless, using dyes to label therapeutics has several drawbacks, including failures to label fluorescent dyes, alteration of properties of drugs, and the possibility that the dyes may detach or disintegrate [16]. It also holds for the widely used methods to evaluate cell therapeutic agents by using tracer labeling.

This study introduced the spatial mapping of exogenous nucleic acids by using ST in a label-free manner. Previous studies have attempted to map RNA transcripts from different species in applications such as xenografts [17], host-microbiome mapping [18], host-virus mapping [19], and the identification of engineered oligonucleotides [20]. Although these studies demonstrated successful analysis on mixed organisms, they did not employ this separated mapping technique to identify the spatial distribution of cell therapeutics in tissues or estimate their mode of action. In addition, our method can be applied to cell therapeutic agents with the same origin as the host when introducing transfection of genes that do not exist in the host [21]. Our proposed approach has the potential to expand the application of ST for spatial analysis of therapeutics containing exogenous nucleic acids. By using this approach, we can gain a more comprehensive understanding of the mechanisms underlying the therapeutic effects of these agents, which can lead to critical insights for optimizing cell therapy treatments for various diseases.

Apart from simple distribution analysis, spatial transcriptomics analysis of stem cell treatment in our study can reveal the transcriptome-level effects on lung fibrosis tissue. The injected human stem cells showed the upregulation of several genes of hemoglobin genes including *Hba-a1, Hba-a2, Hbb-bs*, and *Hbb-bt*, which were upregulated in ‘*Exp’* when compared with ‘*Con*’ group (**Supplementary Figure 4, 6**). In addition, the ‘*Exp*’ group showed downregulated ‘*collagen-containing extracelluar matrix*’ and ‘*humoral immune response*’ genes (*e*.*g*., *Bpifa1* [22]) when compared with ‘*Con*’ (**Figure 2, Supplementary Figure 4-5**). The molecular changes observed in the ‘*Exp*’ group were similar to those identified when comparing the ‘*Nor*’ group with the ‘*Con*’ group (*e*.*g*., *Hba-a1, Hba-a2, Hbb-bs*, and *Hbb-bt*) (**Supplementary Figure 7-8**), which supports the molecular changes were presumed to be the effect of the injected human cells. The comparison of groups at the spot level has limitations because the analysis is only conducted at a single time point after the administration of stem cells. To understand the mechanisms of stem cells at the whole transcriptome level, further experiments in addition to the in-tissue level distribution analysis is needed. This requires using replicate samples and multiple time points.

Several challenges should be noted to use spatial analysis of administered cells using ST. The analytic process presented here was dependent on less significant false positive human transcripts in ‘*Nor*’ and ‘*Con*’. The occurrence of false positive human transcripts came from the homology between human and mouse. Also, false negative human transcripts should be addressed. In the top 20 spatially associated genes with human transcripts in the ‘*Exp*’ sample, *VIM* and *Vim* appeared (**Figure 3a**). Considering *VIM* is a mesenchymal stem cell-associated gene [23], it is highly probable that *Vim* counts were falsely classified mouse transcripts. To resolve the problems presented here, a possible solution is to adopt long-read sequencing to fully manipulate the genetic sequence differences. Additionally, another possible method is to develop computational algorithms that can manipulate spatial proximity [24], histological morphology [25], previous datasets [11, 26] or other domain knowledge to credibly eliminate or modify false transcripts.

Our study successfully demonstrated the ability of ST to map administered cell therapeutics without the need for fluorescent labeling. As a key result, we identified genes associated with human RNA transcripts and showed the spatial distribution of injected human stem cells in lung fibrosis tissue, even though further experiments are needed to fully understand the mechanisms of stem cells in terms of cellular interaction in the target tissue at the whole transcriptome level. Overall, the study highlights the potential of ST for spatial analysis of therapeutics containing exogenous nucleic acids and the need for further research to optimize this approach.

## Methods

### Animal experiments and tissue acquisition

The animal experiments for this study were approved by Seoul National University Bundang Hospital with the IACUC approval code of BA-2211-355-002. Normal bone marrow derived hMSC was prepared from Lonza™. Then, three C57BL/6 male mice (9 weeks old) were prepared: a normal mouse (‘*Nor*’), a mouse with lung fibrosis as a control (‘*Con’*), and a mouse with lung fibrosis and hMSC injection through mouse tail veins as an experimental group (‘*Exp*’). Lung fibrosis was induced in both the control (*‘Con’*) and experimental (*‘Exp’*) groups by administering bleomycin for a period of 3 weeks before sacrifice. Then, optimal cutting temperature (OCT) blocks (Scigen 4586, USA) were made according to the Visium Spatial Protocols – Tissue Preparation Guide (Document CG000240). The OCT block for ‘*Exp*’ was made 6 hr after human stem cell injection. After that, two tissue slices were prepared for each OCT mold of a group to acquire an H&E-stained tissue slice and a fresh frozen tissue slice for Visium ST library.

### ST library acquisition

The tissue sections were fixed, stained, and permeabilized, consulting the Visium Spatial Protocols – Spatial Gene Expression Imaging Guide (Document CG000241), along with tissue optimization (TO) steps. The mRNAs present in the tissues were captured through poly-A tails, and subsequent cDNAs were barcoded and amplified via poly chain reaction (PCR) to obtain enough cDNAs for reconstructing libraries. Quantitative PCR (qPCR) and the Agilent Technologies 4200 TapeStation were used to measure and assess the quality of the libraries, respectively. Finally, the libraries were sequenced using Illumina HiSeq platform, following the instructions provided in the user guide.

In addition, SpaceRanger (ver. 2.0.1) *mkref* was used to merge the representative human reference, *GRCh38*, and the representative mouse reference, *mm10*, followed by the execution of SpaceRanger *count* for each sample to produce the processed ST library.

### Acquisition of DEGs related to hMSC distribution

Data integration was performed with *FindIntegrationAnchors* and *IntegrateData* in Seurat (ver. 4.3.0). Then, spatial clustering analysis was performed after eliminating human genes from the integrated Seurat object. The elimination process was thought to be meaningful not to make biases toward ‘*Exp*’ which only contained human transcripts. However, the elimination process was not made in the other analyses. The top 20 spatially enriched genes (adjusted p value < 0.05; log FC ordered) in a specific cluster were obtained from the Wilcoxon Rank Sum test by using *FindAllMarkers* of Seurat and explored in gene ontology (GO) plot with clusterProfiler::*enrichGO* (ver. 4.6.2) in R.

To perform comparison analysis among samples, differentially expressed genes (DEGs) were acquired from the Wilcoxon Rank Sum test by using *FindAllMakers* of Seurat. Then, top 20 DEGs (adjusted p value < 0.05; log FC ordered) were explored in GO analysis.

To obtain spatially associated genes with the %human among total transcripts, the spearman correlation coefficient between %human and the scaled expression of each gene was calculated in the ‘*Exp*’ sample. After that, top 20 spatially associated genes (adjusted p value < 0.05; spearman correlation ordered) were explored in GO analysis.

### CellDART

To identify spatial distributions of cell types, cell type inference by domain adaptation of single-cell and spatial transcriptomic data (CellDART) was performed [11]. The single cell RNA sequencing (scRNA-seq) reference was created by utilizing a mouse lung scRNA-seq reference (GSE124872). Only focusing on the ‘*Exp*’ sample, CellDART was performed to identify mouse cell types associated with human transcripts.

### Statistics

R (ver 4.0.5) and Python (ver 3.7.12) were used as programming languages. In addition, *Seurat* (ver 4.3.0), *scanpy* (ver 1.9.1), and *SpaceRanger* (ver 2.0.1) were used. For reference for *SpaceRanger*, GRCh38 (*Homo sapiens*) and mm10 (*Mus musculus*) were applied. Differentially expressed genes (DEGs) were explored by sorting against fold change (FC) values for all genes with adjusted p value less than 0.05. When drawing gene ontology (GO) plots, 20 host mouse genes were selected and analyzed unless otherwise specified. The parameters were set to default unless otherwise specified in **Table 1**.

**Table 1.** Specified parameter values for analyses.

| Function | Parameter | Value |
| --- | --- | --- |
| ***Seurat in R*** |  |  |
| ***FindIntegrationAnchors*** | *dims* | 1:50 |
| ***IntegrateData*** | *dims* | 1:50 |
| ***FindNeighbors*** | *dims* | 1:30 |
| ***FindClusters*** | *resolution* | 0.2 |
| ***RunUMAP*** | *dims* | 1:30 |
| ***scanpy in Python*** |  |  |
| ***scanpy.tl.pca*** | *svd_solver* | arpack |
| ***scanpy.pp.neighbors*** | *n_neighbors* | 10 |
| ***scanpy.pp.neighbors*** | *n_pcs* | 40 |
| ***scanpy.tl.leiden*** | *resolution* | 0.5 |
| ***scanpy.pl.rank_genes_groups*** | *n_genes* | 40 |

## Authors Contribution

Hongyoon Choi and Hyung-Jun Im designed this project. Preparation of mouse lung fibrosis model, acquisition of lung tissues and tissue optimization were done by Jae Eun Lee, In Ho Song, and Hyun Soo Park. The library preparation for spatial transcriptomics were performed by Jae Eun Lee. Jeongbin Park and Dongjoo Lee conducted major bioinformatics analyses. D.S.L., H.C., and H.J.I. performed data analysis and interpretation. The manuscript was written by J.P., H.C., and H.J.I. and all authors have contributed to the completion of the manuscript.

## Conflict of Interest

Jeongbin Park, Dongjoo Lee, and Jae Eun Lee are researchers in Portrai, Inc. Daeseung Lee, Hongyoon Choi, and Hyung-Jum Im are the co-founders of Portrai, Inc. In Ho Song is a researcher and Hyun Soo Park is a founder of Molim, Inc. Hyung-Jun Im is a consultant of Cellbion. Schematic illustrations were created with BioRender. Otherwise, there is no competing financial interest.

