## Supplementary Figures 1-8 for "A Spatial Transcriptomics based Label-Free Method for Assessment of Human Stem Cell Distribution and Effects in a Mouse Model of Lung Fibrosis"

**Supplementary Figure Legends**

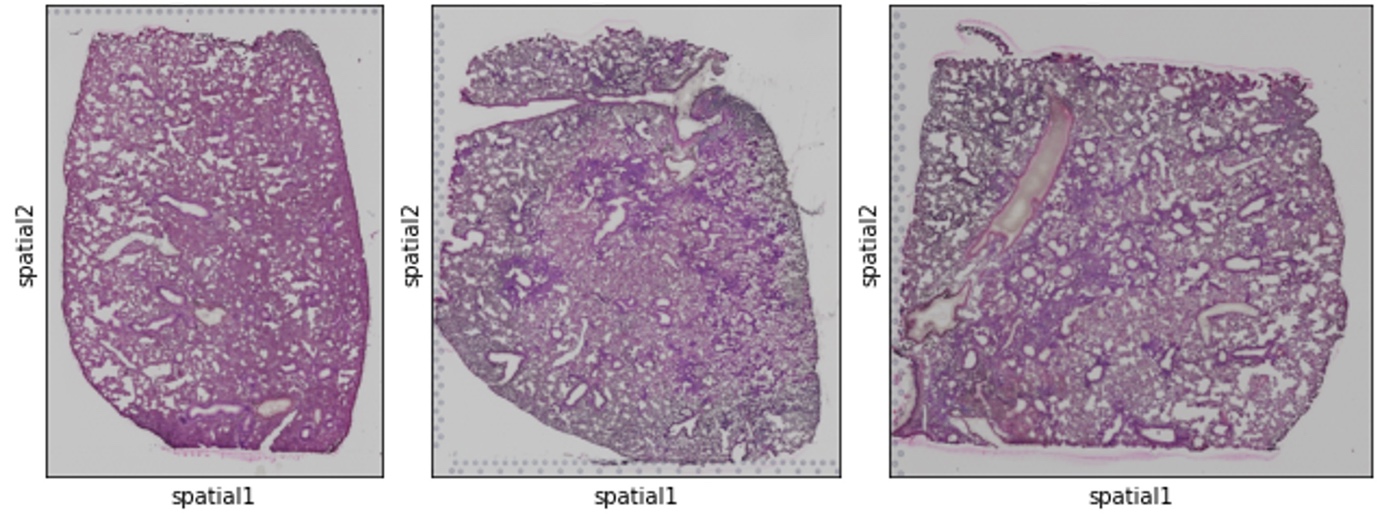

**Supplementary Figure 1** | Histological images for three samples. The images for normal, fibrosis model as a control, and fibrosis model treated with hMSCs were represented from left to right in order.

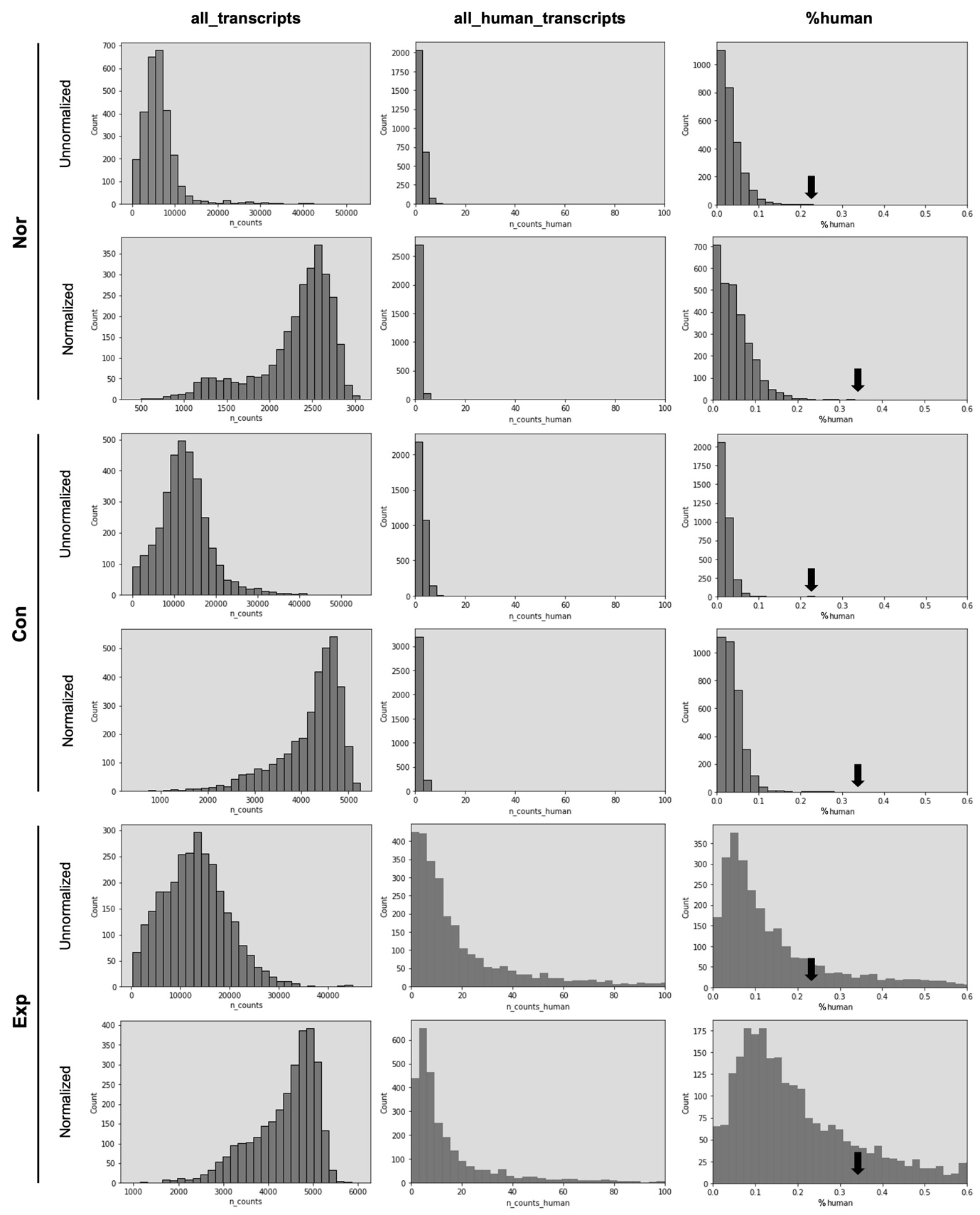

**Supplementary Figure 2** | Histograms of ‘*all_transcripts*’, ‘*all_human_transcripts*’, and %human according to sample and normalization. Here, ‘*Unnormalized*’ and ‘*Normalized*’ covered raw and normalized gene counts, respectively. The number of human transcripts was represented on a scale of 0-100 transcript number and the %human was shown on 0-0.6% scale. The black arrows in the third column indicated the average plus 10 standard deviations of %human of the ‘*Con*’ sample (*i.e.*, 0.23 for the unnormalized and 0.34 for the normalized).

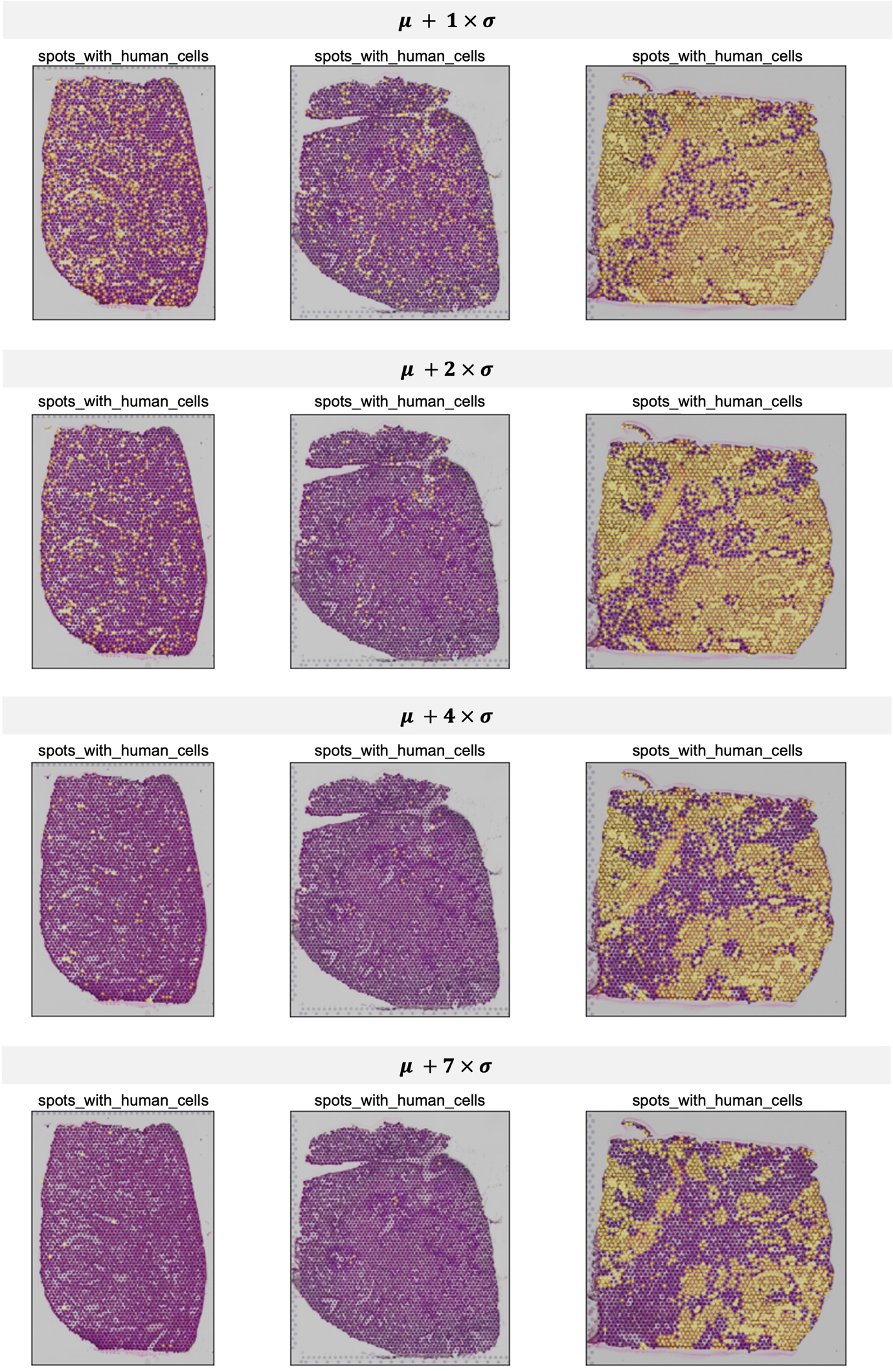

**Supplementary Figure 3** | The representation of ‘*spots_with_human_cells*’ variable that is a binary indicator that is yellow only when the human transcripts on a spot exceed a threshold. The threshold was defined as the average plus 1, 2, 4, or 7 standard deviations of %human of the ‘*Con*’ sample.

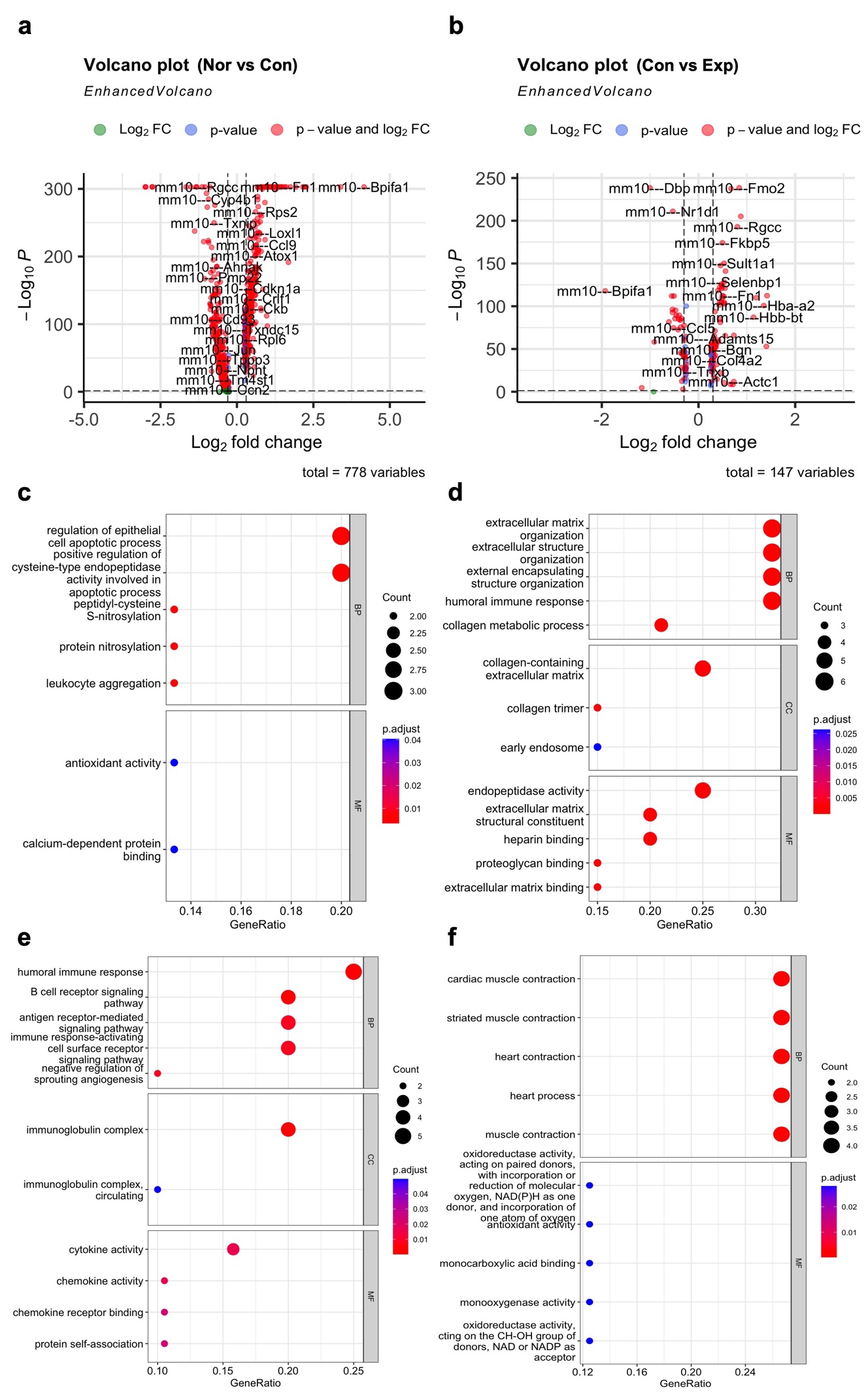

**Supplementary Figure 4 |** Comparison of ‘*Con*’ to the other samples. **a**, A volcano plot comparing ‘*Nor*’ and ‘*Con*’. ‘*Log_2_ fold change*’ > 0 indicated genes enriched in the ‘*Con*’ sample. **b**, A volcano plot comparing ‘*Con*’ and ‘*Exp*’. ‘*Log_2_ fold change*’ > 0 indicated genes enriched in the ‘*Exp*’ sample. Genes annotated with ‘*mm10*’ and ‘*GRCh38*’ were genes from mice and humans, respectively. **c**, GO plots for top 20 DEGs (adjusted p-value < 0.05; log FC ordered) enriched in ‘*Nor*’ compared to ‘*Con*’. **d**, GO plots for top 20 DEGs (adjusted p-value < 0.05; log FC ordered) enriched in ‘*Con*’ compared to ‘*Nor*’. **e**, GO plots for top 20 DEGs (adjusted p-value < 0.05; log FC ordered) enriched in ‘*Con*’ compared to ‘*Exp*’. **f**, GO plots for top 20 DEGs (adjusted p-value < 0.05; log FC ordered) enriched in ‘*Exp*’ compared to ‘*Con*’.

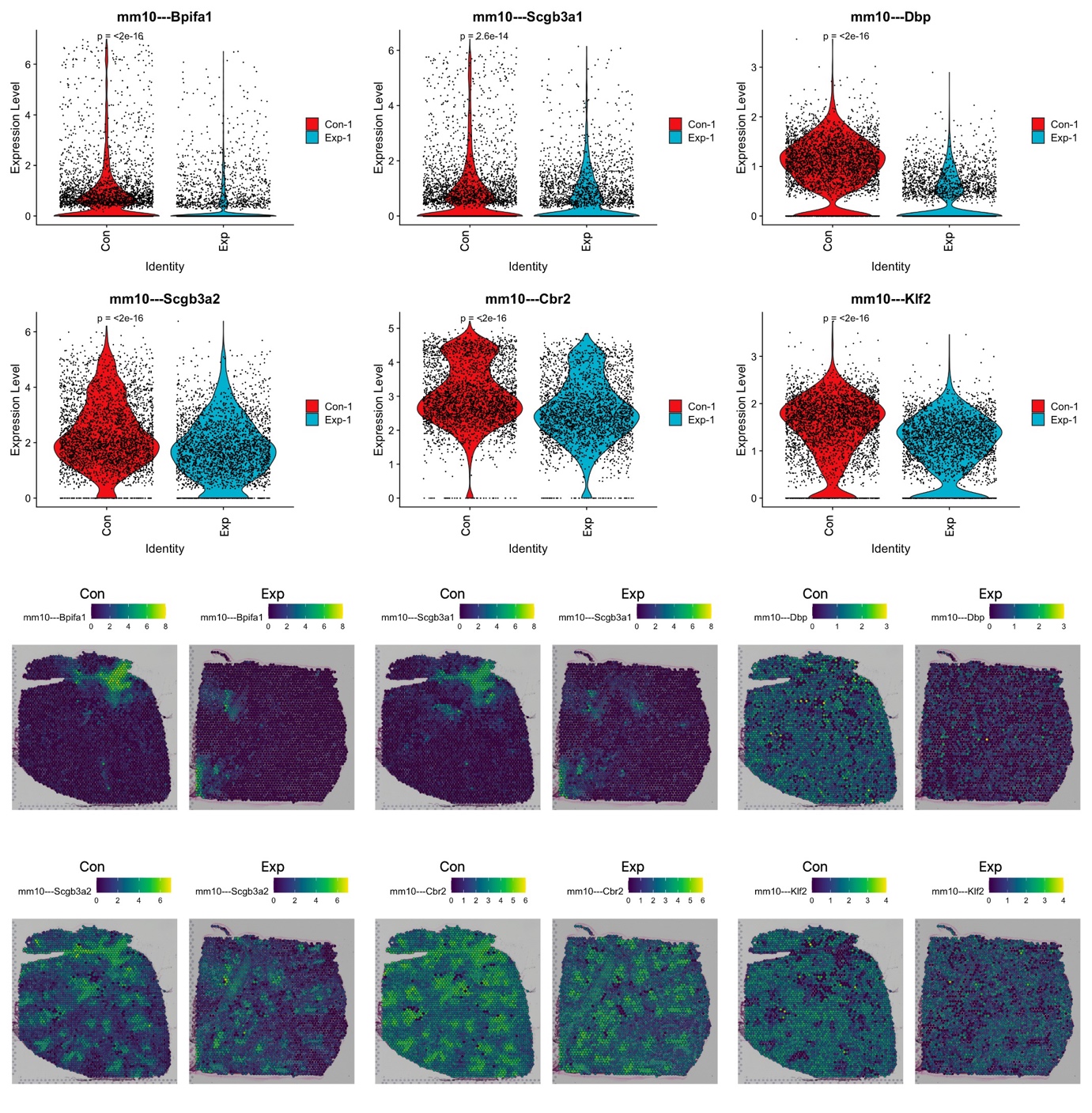

**Supplementary Figure 5** | Top 6 DEGs (adjusted p-value < 0.05; log FC ordered) of ‘*Con*’ compared to ‘*Exp*’.

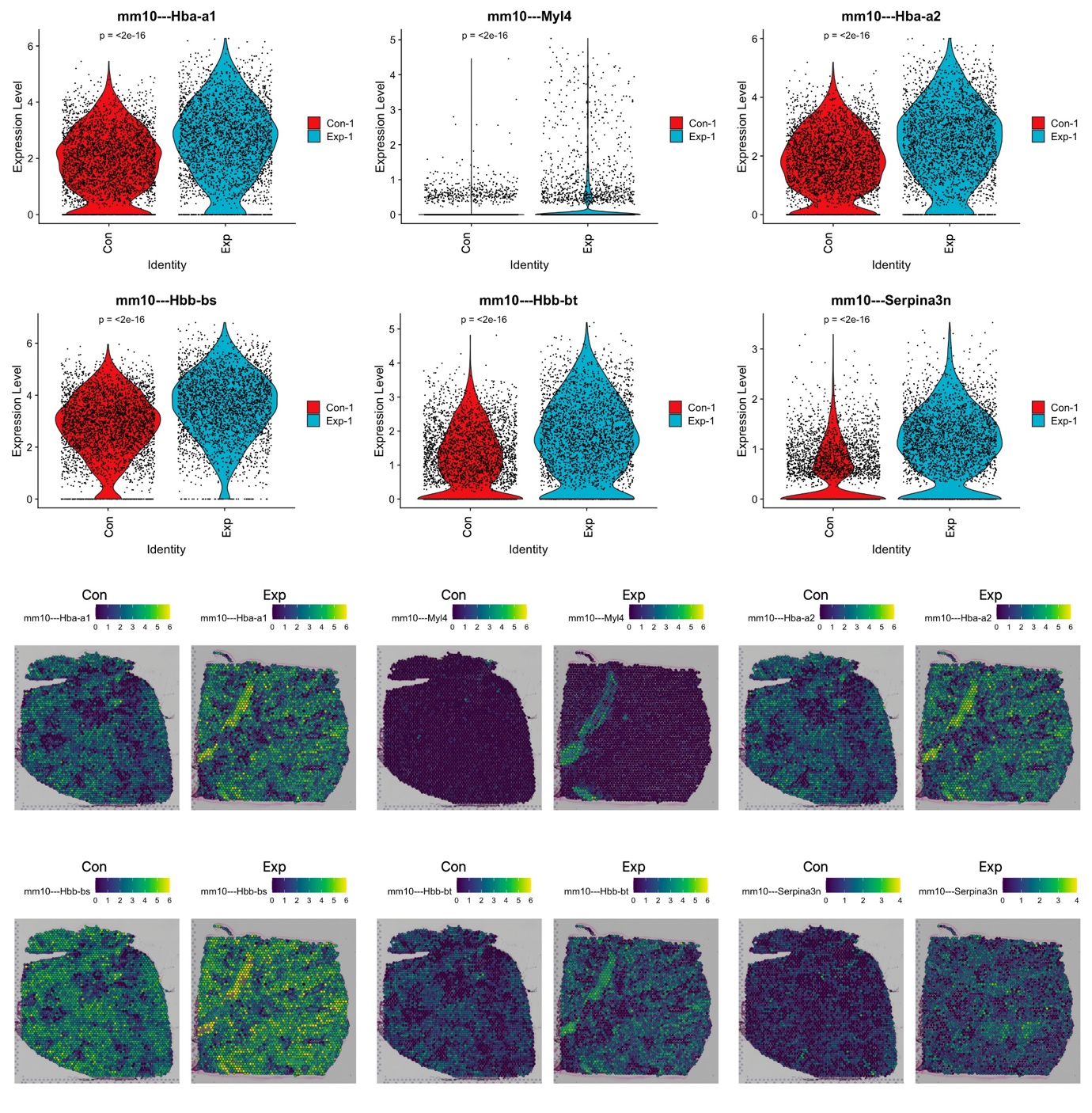

**Supplementary Figure 6** | Top 6 DEGs (adjusted p-value < 0.05; log FC ordered) of ‘*Exp*’ compared to ‘*Con*’.

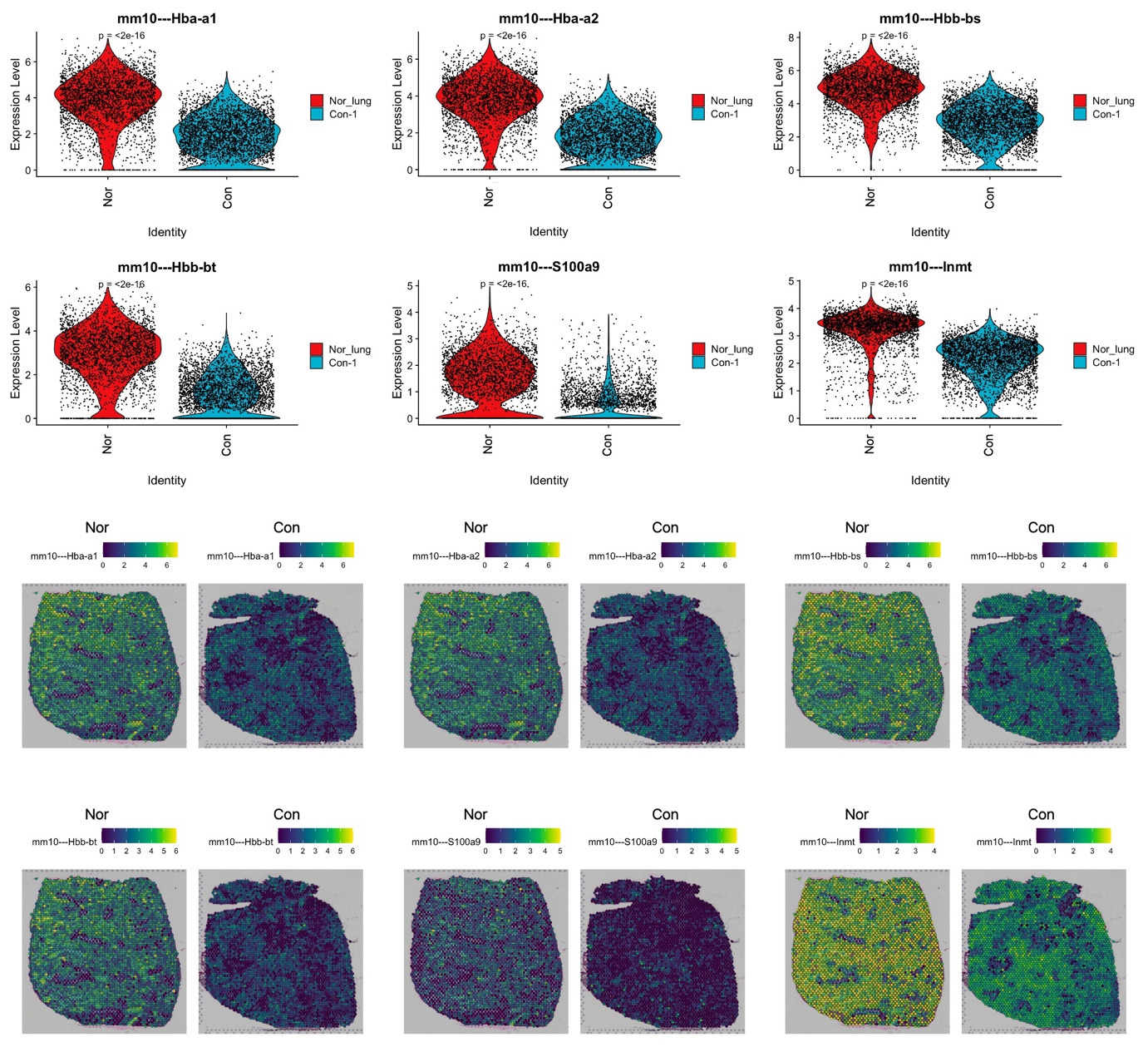

**Supplementary Figure 7** | Top 6 DEGs (adjusted p-value < 0.05; log FC ordered) of ‘*Nor*’ compared to ‘*Con*’.

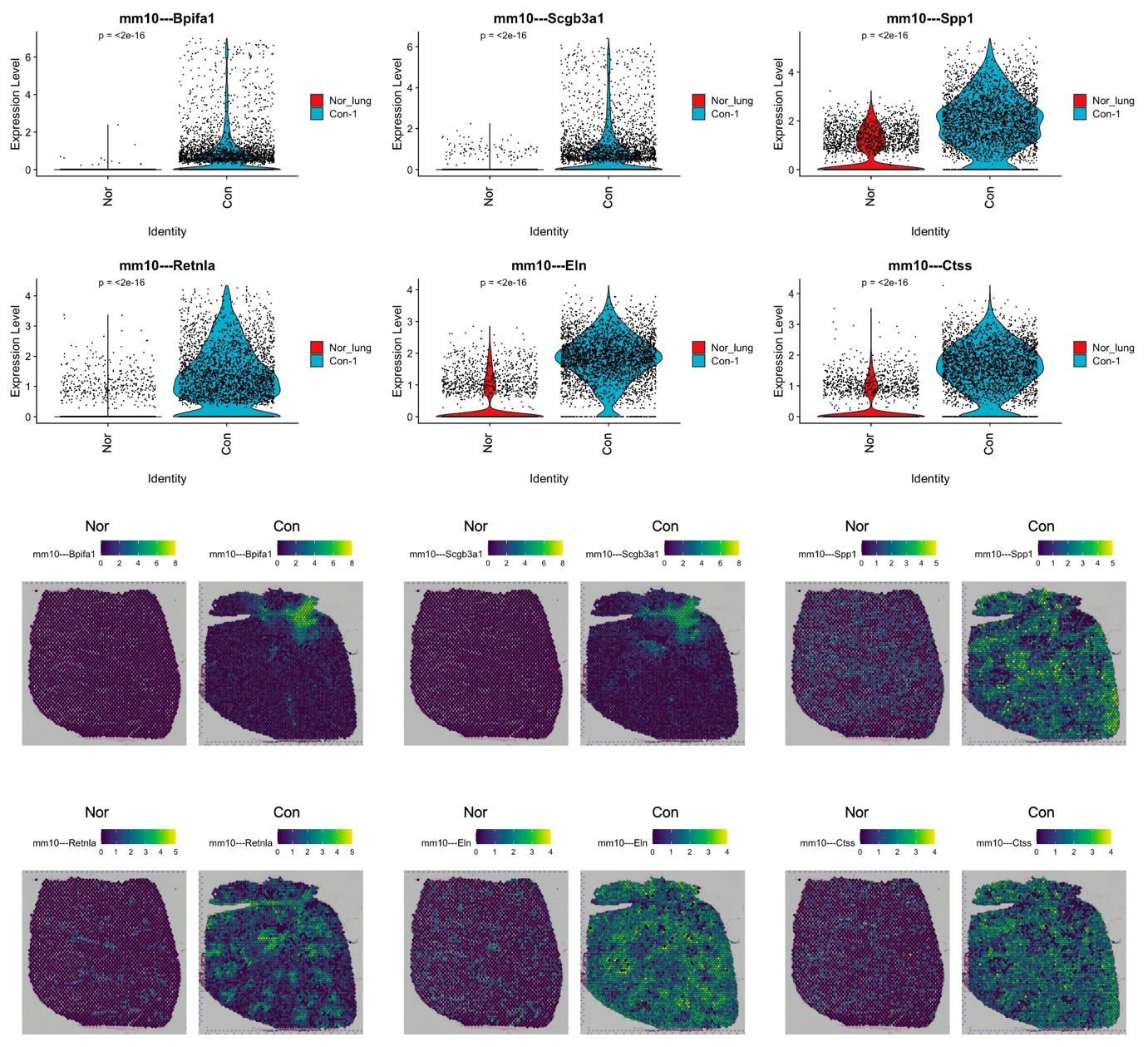

**Supplementary Figure 8** | Top 6 DEGs (adjusted p-value < 0.05; log FC ordered) of ‘*Con*’ compared to ‘*Nor*’.
